# “Unexpected mutations after CRISPR-Cas9 editing *in vivo*” are most likely pre-existing sequence variants and not nuclease-induced mutations

**DOI:** 10.1101/159707

**Authors:** Caleb A. Lareau, Kendell Clement, Jonathan Y. Hsu, Vikram Pattanayak, J. Keith Joung, Martin J. Aryee, Luca Pinello

**Author notes:** These authors contributed equally to this work (Caleb A. Lareau, Kendell Clement, Jonathan Y. Hsu).

## Abstract

Schaefer *et al*. recently advanced the provocative conclusion that CRISPR-Cas9 nuclease can induce off-target alterations at genomic loci that do not resemble the intended on-target site.^1^ Using high-coverage whole genome sequencing (**WGS**), these authors reported finding SNPs and indels in two CRISPR-Cas9-treated mice that were not present in a single untreated control mouse. On the basis of this association, Schaefer *et al*. concluded that these sequence variants were caused by CRISPR-Cas9. This new proposed CRISPR-Cas9 off-target activity runs contrary to previously published work^2–8^ and, if the authors are correct, could have profound implications for research and therapeutic applications. Here, we demonstrate that the simplest interpretation of Schaefer *et al.’s* data is that the two CRISPR-Cas9-treated mice are actually more closely related genetically to each other than to the control mouse. This strongly suggests that the so-called “unexpected mutations” simply represent SNPs and indels shared in common by these mice prior to nuclease treatment. In addition, given the genomic and sequence distribution profiles of these variants, we show that it is challenging to explain how CRISPR-Cas9 might be expected to induce such changes. Finally, we argue that the lack of appropriate controls in Schaefer *et al.’s* experimental design precludes assignment of causality to CRISPR-Cas9. Given these substantial issues, we urge Schaefer *et al*. to revise or re-state the original conclusions of their published work so as to avoid leaving misleading and unsupported statements to persist in the literature.

---

The conclusion that the sequence variants shared by the genome-edited F03 and F05 mice (and not found in the control untreated FVB mouse) are caused by CRISPR-Cas9 ***critically depends*** upon the assumption that all of these mice were initially genetically identical. Indeed, the authors have repeatedly relied on the assertion that the mice used are from a highly inbred strain. If this clonality assumption were true, one would expect that all three mice should be nearly identical for common variants found in dbSNP^9^ (a database of known SNPs and indels that Schaefer *et al*. used to filter and exclude potential false positive mutations that may be attributed to CRISPR-Cas9 activity), with no two mice sharing genotypes at any loci that are not the same in the third mouse (a hypothetical result represented in Fig. 1a). However, after genotyping these mice with GATK best practices, we identified a total of 31,079 high-quality variants at dbSNP loci that were concordant in two mice but distinct from the third when examining all possible pairwise combinations (Fig. 1b; **Supplementary Note 1**). We also note that many (33-46%) of the high-confidence genotyped variants in each mouse are heterozygous (**Supplementary Table 1**), which the authors have argued should not exist in highly inbred mice.^10^ These observations clearly demonstrate that the three mice are neither clonal nor completely isogenic.

**Figure 1.**
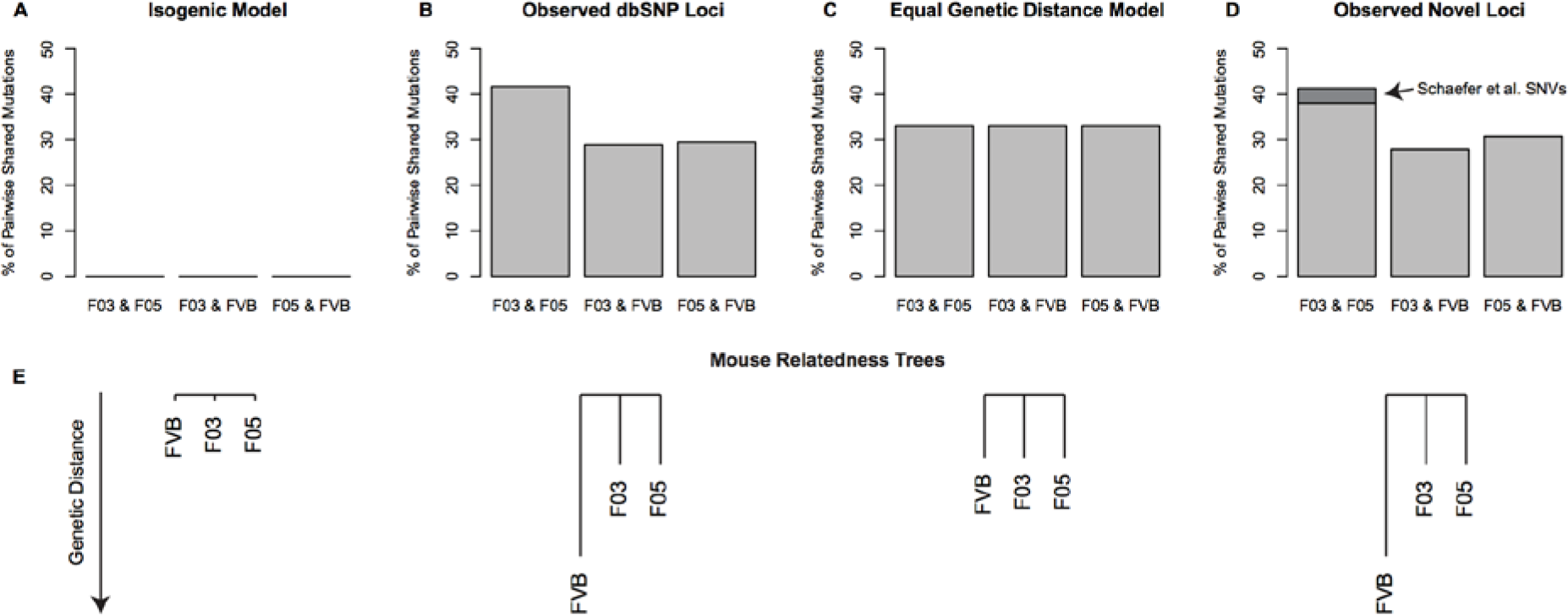
Measures of genetic relatedness in the F03, F05 and FVB mice. Schaefer *et al.’s* experimental model to examine mutations attributable to Cas9 treatment relies on the assumption that mice are isogenic, meaning no private mutations are observed within or shared between mice as depicted in (A). However, after genotyping these mice using GATK best practices, we observe a significant departure from this model, suggesting that the F03 and F05 mice are more genetically related at common variants (B; n = 31,079). As an isogenic system is practically impossible, the selection of littermate assumes the number of loci with shared genotypes is nearly identical for all mice, leading to equivalent genetic distances separating them as shown in (C). This representation of an equal genetic model demonstrates a clear departure from the observed data at common variants and other non-dbSNP loci that we term “novel variants” (D; n = 38,981). Additionally, the variants previously reported by Schaefer *et al.* (dark gray, D) represent a small subset of the genotypes common to F03 and F05 but distinct from FVB at non-dbSNP sites. The observed ratios in B and D cannot be distinguished from each other (p = 0.304; two-sided Fisher’s Exact Test), but each represent a significant departure (p < 2.2 10^−16^; Chi-Squared Test) from the equal genetic distance model (C) required to attribute differential SNVs to Cas9 activity. Panel (E) provides a graphical summary of these models of genetic relatedness using under the two hypothetical models and the two sets of observed variants.

Because perfectly isogenic mice are essentially impossible to obtain, we suggest that an equal distance model that allows rare and private mutations may be more realistic and more appropriate (a hypothetical result is illustrated **in** Fig. 1c). Even considering this relaxed model, our analysis of Schaefer *et al.’s* WGS data clearly reveals several ways in which the F03 and F05 mice appear to be genetically more closely related to each other than to the control FVB mouse: First, the percentage of dbSNP loci (variants that were filtered out by Schaefer *et al.’s* workflow^1^) shared between F03 and F05 compared with FVB is higher (41.7%) than those found with the two other pairwise comparisons among the three mice (28.95% and 29.4%; Fig. 1b). The skew in this distribution differs significantly (p < 2.2 × 10^−16^; Chi-Squared Test) from what is expected if the three mice were equally genetically distant from one another (compare **Figs. 1b** and **1c**). Second, in comparing the distribution of known dbSNP variants to “novel” sequence variants (including the ones attributed to CRISPR-Cas9 by Schaefer *et al.*; see **Supplementary Note 1**), we could not reject (p = 0.304; two-sided Fisher’s Exact Test) the null hypothesis that the rates of shared pairwise genotypes with these “novel” sequence variants (Fig. 1d) differed from the distribution of dbSNP variants (compare **Figs. 1b** and **1d**). Third, we constructed genetic relatedness trees (Fig. 1e; **Supplementary Note 2**) from all the sequence variants, which showed that the F03 and F05 samples were more closely related to each other than to the FVB control. These distances differ from the hypothetical case in which the mice are equally genetically distant from one another. Collectively, these analyses show that the variants at non-dbSNP loci (including those attributed to CRISPR activity) do not co-occur more frequently in the F03 and F05 mice than one would expect. Instead, we find that the CRISPR-treated mice are genetically more similar to each other than to the control FVB mouse.

Even if one were to assume that the variants in question were induced by CRISPR-Cas9, it is extremely challenging to reconcile the new mutagenic off-target activity proposed by Schaefer *et al*. with our current extensive understanding of how this nuclease functions. For example, as the authors themselves correctly note, no DNA sequences resembling the on-target site can be found near the sequence variants they attribute to CRISPR-Cas9, a result we re-validated with our own analysis of their WGS data (**Supplementary Fig. 2; Supplementary Note 3**). This finding is puzzling given the high concordance in the genomic locations (>1,300 sites reported by Schaefer *et al*. and ~30,000 discovered using GATK best practices) of these sequence variants in the F03 and F05 mice. If these alterations are somehow being caused by CRISPR-Cas9 then this suggests that the nuclease must be directing alterations consistently to the same genomic addresses by recognizing some type of DNA sequence common to these loci. However, our attempt to find a consensus DNA sequence at or near the locations of these sequence variants failed to reveal any consistent motif (**Supplementary Fig. 3; Supplementary Note 4)**, a result that makes it hard to envision any reasonable mechanism for how CRISPR-Cas9 could direct alterations to the same genomic loci in the two mice. Furthermore, it is very striking how consistent the indels purportedly induced by CRISPR-Cas9 are between the two mice (**Supplementary Fig. 4**). In particular, because CRISPR-Cas9 is known to induce indels at target sites that are heterogeneous in both length and positioning relative to the DNA break, it would be highly unlikely that, if the nuclease were inducing indel mutations in the two mice, that these changes would be exactly the same length and result in the same altered sequence at a large number of loci. Indeed, our modeling using empirical data from actual CRISPR-Cas9-modified sites^11^ estimates that the probability of 118 indels being exactly the same length in the F03 and F05 mice is less than 1 in 10^12^ under the most generous assumptions (binomial distribution; **Supplementary Note 5**). Overall, it is very difficult to envision or articulate how CRISPR-Cas9 would mechanistically induce the alterations ascribed to the nuclease by Schaefer *et al*.

Given these various issues, how then might the sequence differences attributed to CRISPR-Cas9 by Schaefer *et al*. best be explained? Based on our analysis of their reported data, the simplest explanation is that the F03 and F05 mice are more closely related genetically than the control FVB mouse. That is, the CRISPR-treated embryos most likely already harbored these private SNPs and indels prior to nuclease treatment whereas the control mouse did not. Further support for this idea comes from our finding that the putative variants attributed to CRISPR-Cas9 by Schaefer *et al*. were significantly closer in distance to common variants in dbSNP shared between F03 and F05 but not in FVB (p < 2 × 10^−19^, two-sided Fisher’s Exact Test; **Supplementary Note 6**), strongly suggesting that those variants were likely already present and co-inherited by F03 and F05 prior to nuclease treatment (**Supplementary Fig. 6**). Furthermore, the observed heterozygous and homozygous frequencies of these sequence variants in the mosaic F03 and F05 animals are also entirely consistent with this hypothesis. Our proposed alternative explanation for these “unexpected mutations” also obviates the need to invoke a new highly efficient activity of CRISPR-Cas9 that has neither been seen to date nor can be readily understood based on previously reported insights into how these nucleases function.

Finally, we note that Schaefer *et al.’s* incorrect interpretation of their data could likely have been avoided by performing appropriate controls. Although the authors clearly identified an association between the sequence variant landscape and Cas9 treatment, this observation alone does not prove causality without proper consideration of potential confounding factors, in particular the genetic background of these mice. For example, using a permutation of the authors’ own WGS analysis framework, we observed an equally high percentage of heterozygous variants in the control mouse that are not present in the two nuclease-treated mice (**Supplementary Table 1**), but we would certainly not attribute these as mutations induced by the lack of CRISPR-Cas9 treatment in the control mouse. Had Schaefer *et al.* performed a few key control experiments (e.g., using only Cas9 with no guide RNA), these experiments would likely have yielded results demonstrating that CRISPR-Cas9 is not responsible for the sequence differences in question.

In light of all of the above (and critiques published by others^12,13^ while we were finalizing this manuscript), we strongly encourage the authors to restate (or at least temper) the title and conclusions of their original paper or provide properly controlled experiments that can adequately support their claims. Not doing so does a disservice to the field and leaves the misleading impression that the strong statements and recommendations found in their paper are adequately supported by the data presented.

**Data availability statement.** Sequencing data as part of the original study was accessed from available at SRA accessions SRR5450996-SRR5450998. Source data for Figure 1 is available as part of our online resource: http://aryeelab.org/crispr_mutation_reanalysis

**Code availability statement.** Code and data to reproduce our analysis can be visualized and download from our online repository: http://aryeelab.org/crispr_mutation_reanalysis

*Note: Any Supplementary Information and Source Data files are available in the online version of the paper.*

## Acknowledgments

This work was supported by the National Science Foundation (DGE1144152 to C.A.L), National Institutes of Health (R35 GM118158 to J.K.J. and R00 HG008399 to L.P.), the Defense Advanced Research Projects Agency (HR0011-17-2-0042 to J.K.J. and L.P.), and the Desmond and Ann Heathwood Massachusetts General Hospital Research Scholar Award (J.K.J.). We thank Benjamin Kleinstiver, Jose Malagon Lopez and Alexander Anthony for helpful discussions.

